# Elevated temperatures drive the evolution of armor loss in the threespine stickleback *Gasterosteus aculeatus*

**DOI:** 10.1101/2020.10.29.361568

**Authors:** Carl Smith, Grzegorz Zięba, Mirosław Przybylski

## Abstract

While there is evidence of genetic and phenotypic responses to climate change, few studies have demonstrated change in functional traits with a known genetic basis. Here we present evidence for an evolutionary adaptive response to elevated temperatures in freshwater populations of the threespine stickleback (*Gasterosteus aculeatus*). Using a unique set of historical data and museum specimens, in combination with contemporary samples, we fitted a Bayesian spatial model to identify a population-level decline in the number of lateral bony plates, comprising anti-predator armor, in multiple populations of sticklebacks over the last 90 years in Poland. Armor loss was predicted by elevated temperatures and is proposed to be a correlated response to selection for reduced body size. This study demonstrates a change in a functional trait of known genetic basis in response to elevated temperature, and illustrates the utility of the threespine stickleback as a model for measuring the evolutionary and ecological impacts of environmental change across the northern hemisphere.

## Introduction

The central assumption of evolutionary theory is that natural selection drives phenotypic adaptation of populations to local environmental conditions through changes in the genetic composition of the population. These adaptive changes arise from the differential reproductive success of individuals that vary genetically (Darwin 1859; Fisher 1930; Williams 1966). In this context, the capacity of natural populations to respond to rapid climate change is contingent on adaptive change to phenotypic traits that have a genetic basis. While numerous studies demonstrate correlations between temperature and phenotypic traits, many traits are highly plastic and the adaptive value of these changes is unclear (reviewed by Crozier and Hutchings 2014).

To unequivocally show an adaptive evolutionary response to climate change it is necessary to demonstrate consistent and predicted alteration in a functional trait that is under genetic control. A suitable species is one with short generation time, subject to high or consistent selection pressure for a trait under simple genetic control and with standing genetic variation present in populations (Crozier & Hutchings 2014). The threespine stickleback (*Gasterosteus aculeatus*) is a model vertebrate in behavioral and evolutionary biology that possesses these features and is widely distributed across the northern hemisphere. The species occupies a wide range of environments and shows a high degree of phenotypic variability over small spatial scales (Bell & Foster 1994; Wootton 1976, 1984, 2009; Des Roches et al. 2019; Smith et al. 2020a). Marine populations of the threespine stickleback have repeatedly invaded freshwater habitats. These invasions are characterized by rapid reduction in the extent of anti-predator ‘armor’, comprising lateral bony plates, pelvic girdle and spines, and dorsal spines, as well as other aspects of their biology (Wootton 1976, 2009; Bell & Foster, 1994).

Most freshwater populations of threespine sticklebacks converge on a *low* lateral plate ecomorph, with <10 lateral plates on each side of the anterior portion of the body, along with three dorsal spines, pelvic girdle and a pair of pelvic spines. The separate elements of the armor function in concert, with anterior lateral plates bracing the dorsal and pelvic spines, which thereby resist compression while also limiting ingestion by gape-limited vertebrate predators (Reimchen, 1994). Marine sticklebacks are almost exclusively represented by a *complete* ecomorph, with >30 lateral plates along the entire body. Posterior lateral plates appear to function in preventing skin puncture by toothed predators (Reimchen, 1994). A *partial* ecomorph also occurs, expressing an intermediate and variable number of lateral plates, and is typically encountered in brackish water (Wootton, 1976). Population-level change in the number of lateral plates can be rapid, with a reversal in plate ecomorph dominance in as few as 10 generations (e.g. Bell *et al*., 2004; Kitano *et al*., 2008).

The accepted contemporary model for threespine stickleback lateral plate evolution involves the repeated, independent establishment of freshwater populations from ancestral, panmictic populations of fully-plated marine fish. Multiple genomic regions have been associated with adaptation to fresh water (Jones *et al*., 2012), with the *ectodysplasin A* (EDA) locus specifically implicated in lateral plate evolution. Notably, 70% of variation in plate number and size is associated with variation in EDA (Cresko *et al*., 2004; Colosimo *et al*., 2005). Marine populations are assumed to possess a pool of standing genetic variation, with rare variants of the EDA locus experiencing strong selection once fish enter fresh water, where they rapidly increase in frequency (Barrett, 2010).

A number of hypotheses have been proposed to explain the selective agent responsible for armor loss in freshwaters, including predation, calcium and phosphorus availability, water density, parasitism, competition, body size and swimming performance (Wootton 2009; Myhre & Klepaker, 2009; Barrett, 2010; Smith et al. 2020b). At a large geographic scale, it is also apparent that temperature plays a role in driving variation in lateral plate number (Münzing 1963; Wootton 1976, 1984; Des Roches et al. 2019; Smith et al. 2020b), with well-developed armor associated with low winter temperatures and the converse at higher temperatures. A striking deviation from the typical pattern of lateral plate morph evolution is the widespread occurrence of fully plated sticklebacks in fresh waters in eastern Europe and the east coast of North America, where they are associated with low winter temperatures (Wootton, 1976, 2009; Hagen & Moodie, 1982). The almost ubiquitous occurrence of *complete* ecomorph populations in Poland, in particular, has been the focus of research for many decades (e.g. Piesik 1937; Penczak 1965; Bańbura 1989; Smith et al. 2020b).

The association between environmental temperature and variation in threespine stickleback lateral plate number provides an opportunity to investigate whether large-scale climate trends can drive phenotypic change in lateral plate number among stickleback populations. Their wide distribution, tolerance of a wide range of environmental conditions, and striking phenotypic variability makes the threespine stickleback an ideal model for investigating selective forces underpinning climate change.

In this study, we investigated temporal trends in lateral plate number over a 90-year period across Poland, using a spatially explicit Bayesian model. Given the association between lateral plate number and temperature, we predicted a decrease in the lateral plate number of threespine sticklebacks over the past nine decades, corresponding with increasing environmental temperatures, while controlling for the effects of concomitant changes in body size and spatial dependency.

## Material and Methods

### Fish data

Samples of threespine sticklebacks were collected in 2017 and 2018 from 61 sites across Poland using electrofishing, dipnets and small Seine nets. Collected fish were killed with anesthetic (benzocaine) and fixed in 4% buffered formalin. For each fish, a record was made of standard length (SL, measured from the tip of the snout to the origin of the tail) and total number of lateral plates on the left flank of the body. An additional 15 samples were obtained from the archive of the Museum of the Institute of Zoology of the Polish Academy of Sciences in Warsaw. These samples were collected between 1927 and 1963. Specimens were measured for SL and the total number of lateral plates on the left side of each fish counted. Information on the date and location of collected specimens were available. Finally, we used previously published data from colleagues at the University of Łódź (Penczak 1960, 1962; Bańbura 1988, 1994), that included fish collections conducted between 1947 and 1987 for 78 populations. Thus, data were obtained for 154 Polish populations spanning 91 years (1927-2018). The locations of sampling sites are illustrated in Fig. 1. and summarized in Table S1. Samples were restricted to adult fish >27 mm SL to ensure lateral plate development was complete (Bańbura 1989).

**Fig. 1.**
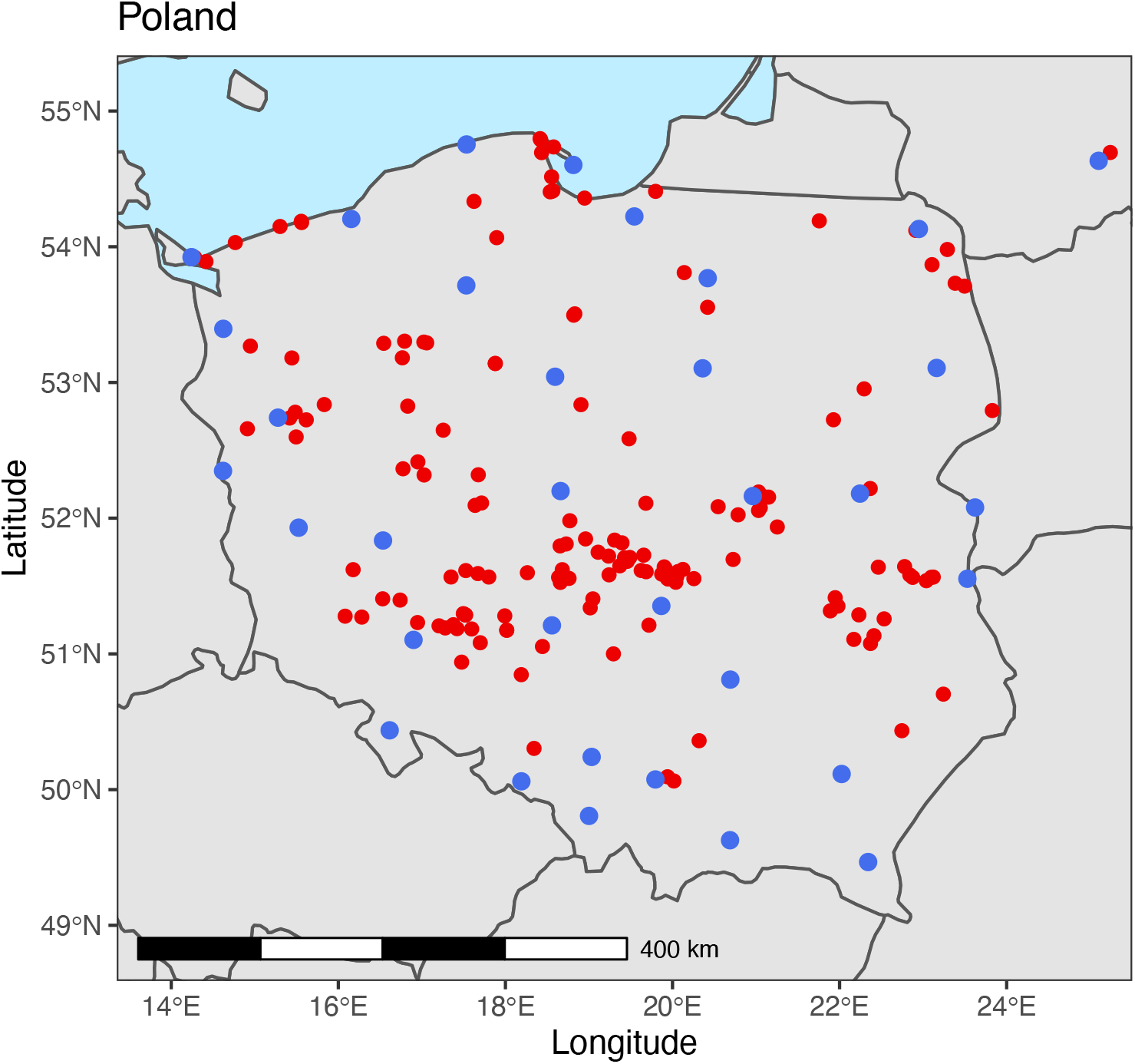
Collection sites of threespine stickleback samples (red dots) and weather stations (blue dots) in Poland. Note that one sample, in the top right of the figure, falls outside the current Polish border. When this sample was collected (September 1928) the collection site was on Polish territory (now in Lithuania).

### Temperature data

Temperature data were compiled from 34 meteorological stations across Poland. Daily air temperature data were available from January 1881 until the present, though not for all weather stations. For each stickleback collection site, the mean of the 10-year antecedent air temperatures (T_10_) from the geographically nearest weather station with a complete time series of data was calculated for each stickleback sampling location. A period of 10 years was used because this is sufficient time for a change in threespine stickleback lateral plate phenotype to evolve following environmental change (Bell et al. 2004). For some sample locations, a complete 10-year time-series of temperature data was not available (data collection was interrupted at some locations during WWII and in its immediate aftermath), in which case an incomplete time series was used. The spatial distribution of weather stations is shown in Fig. 1. Summary data for every sample and the location of the nearest weather station are summarized in Tables S1 and S2.

### Data analysis

Data were modelled using R (version 4.0.2; R Development Core Team 2020) with models fitted in a Bayesian framework using INLA (Integrated Nested Laplace Approximation) with the R-INLA package (Rue et al. 2017). To model the data at the population level, lateral plate count (Gaussian distribution) was regressed against historical temperature data and body size. Data exploration suggested trends in plate number associated with changes in the 10-year antecedent near-surface air temperature (SAT) anomaly over the last nine decades (Fig. 2A), particularly associated with a rapid rise in the European SAT over the last three decades (Fig. 2B). These data included spatial patterns arising from variation in sample locations through time, in combination with spatial variation in SAT among sample locations, resulting in residual spatial autocorrelation (Legendre 1993). To accommodate this potentially confounding structure to the data, we used spatially explicit, Bayesian approximation methods to generate model parameter estimates. R-INLA includes functions to construct Gaussian Markovian random fields (GMRFs) that permit parameter estimation in relation to spatial structure in the data (Lindgren et al., 2011). GMRFs are estimated using Matérn correlation solved with a stochastic partial differential equation (SPDE) on a ‘mesh’, comprising small, non-overlapping triangles that encompass the sampling area (Zuur et al. 2017). Using this approach, we modeled threespine stickleback plate number as a function of body size (SL) and T_10_ from the geographically nearest weather station to each sampling site, while accommodating spatial dependency in the data. We predicted a negative relationship between plate number and T_10_ and a positive relationship with body size (Smith et al. 2020b).

**Fig. 2.**
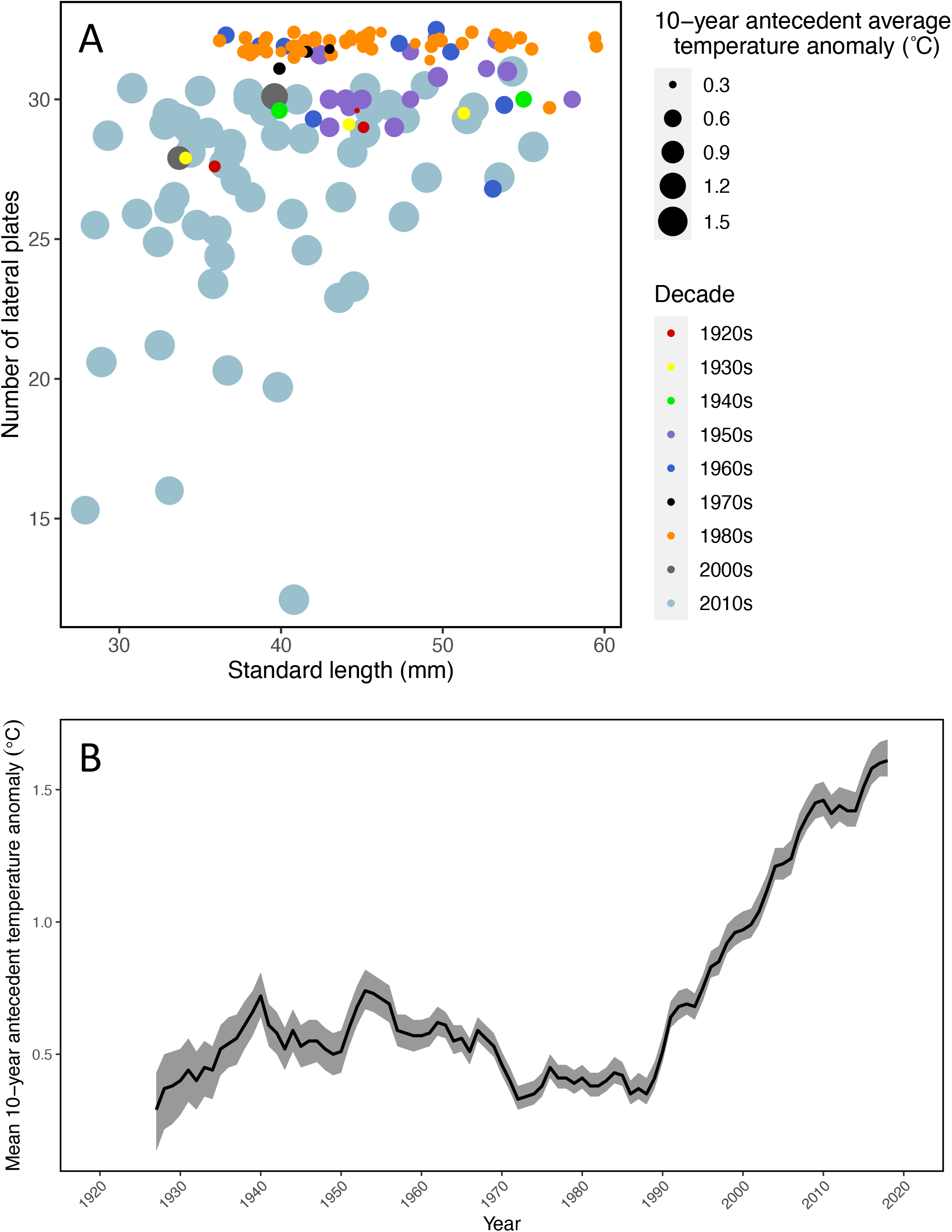
(A) Bubble plot of threespine stickleback lateral plate number against standard length (mm), with size of points relative to mean 10-year antecedent near-surface air temperature (SAT) anomalies over land for Europe (°C) from the CRUTEM4 dataset compiled by the UK Met Office Hadley Centre (www.metoffice.gov.uk/hadobs/hadcrut4). Different colored points represent different decades; (B) Plot of 10-year antecedent SAT anomalies over land for Europe (°C) from the CRUTEM4 dataset by year from 1927 – 2018, the period over which threespine stickleback data were collected and compiled. The black line is the mean SAT and shaded area is the 95% confidence interval.

Seven models were fitted to the data, with SL and T_10_ either included or removed, and with or without the inclusion of a spatially correlated random effect. A baseline model was also included, comprising an intercept and spatially correlated random effect only. Model selection was conducted using the Deviance Information Criterion (DIC) (Spiegelhalter et al. 2002) as a measure of goodness-of-fit, and the Log-Conditional Predictive Ordinate (LCPO) (Roos & Held 2011) as a measure of model predictive quality. The best fitting model, determined by DIC and LCPO, took the form:

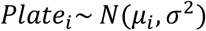

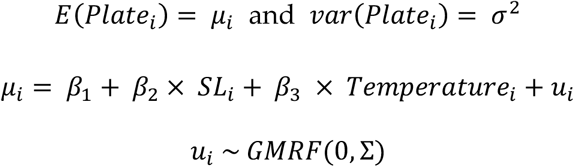

Where *Platei* is the number of lateral plates for the stickleback population at sampling location *i* assuming a Gaussian distribution with mean *μi* and variance *σ2*. The variable *SLi* is a continuous covariate representing population standard length at sampling location *i*, and *Temperaturei* is a continuous covariate representing the mean 10-year antecedent air temperature at location *i*. The term *ui* is a spatially correlated random effect at sampling location *i* with mean 0 and covariance ∑.

## Results

A mesh of 3,874 vertices was generated by constrained refined Delaunay triangulation to encompass all sampling sites in Poland (Fig. 3A). The spatial model was effective in removing residual spatial autocorrelation in the model; model comparison demonstrated that inclusion of a spatial random effect improved model goodness of fit (DIC) and predictive quality (LCPO) in comparison with the same models that did not accommodate spatial dependency (Table 1). Posterior mean values of the GMRF also showed clear spatial patterns (Fig. 3B), as did variance in the posterior values (Fig. 3C). The best-fitting model included fixed effects for SL and T_10_ and a spatially correlated random effect (Table 1).

**Table 1.** Deviance Information Criterion (DIC) and Log-Conditional Predictive Ordinates (LCPO) of fitted models using Integrated Nested Laplace Approximation (INLA) to determine threespine stickleback lateral plate number. SL is stickleback standard length, T_10_ is the average 10-year antecedent air temperature. Models ranked by DIC and LCPO.

| Rank | Model | DIC | LCPO |
| --- | --- | --- | --- |
| 1 | Spatial effect + SL + $T_{10}$ | 693 | 350 |
| 2 | SL + $T_{10}$ | 698 | 351 |
| 3 | Spatial effect + $T_{10}$ | 705 | 357 |
| 4 | $T_{10}$ | 711 | 358 |
| 5 | Spatial effect + SL | 715 | 359 |
| 6 | SL | 719 | 361 |
| 7 | Spatial effect only | 739 | 371 |

**Fig. 3.**
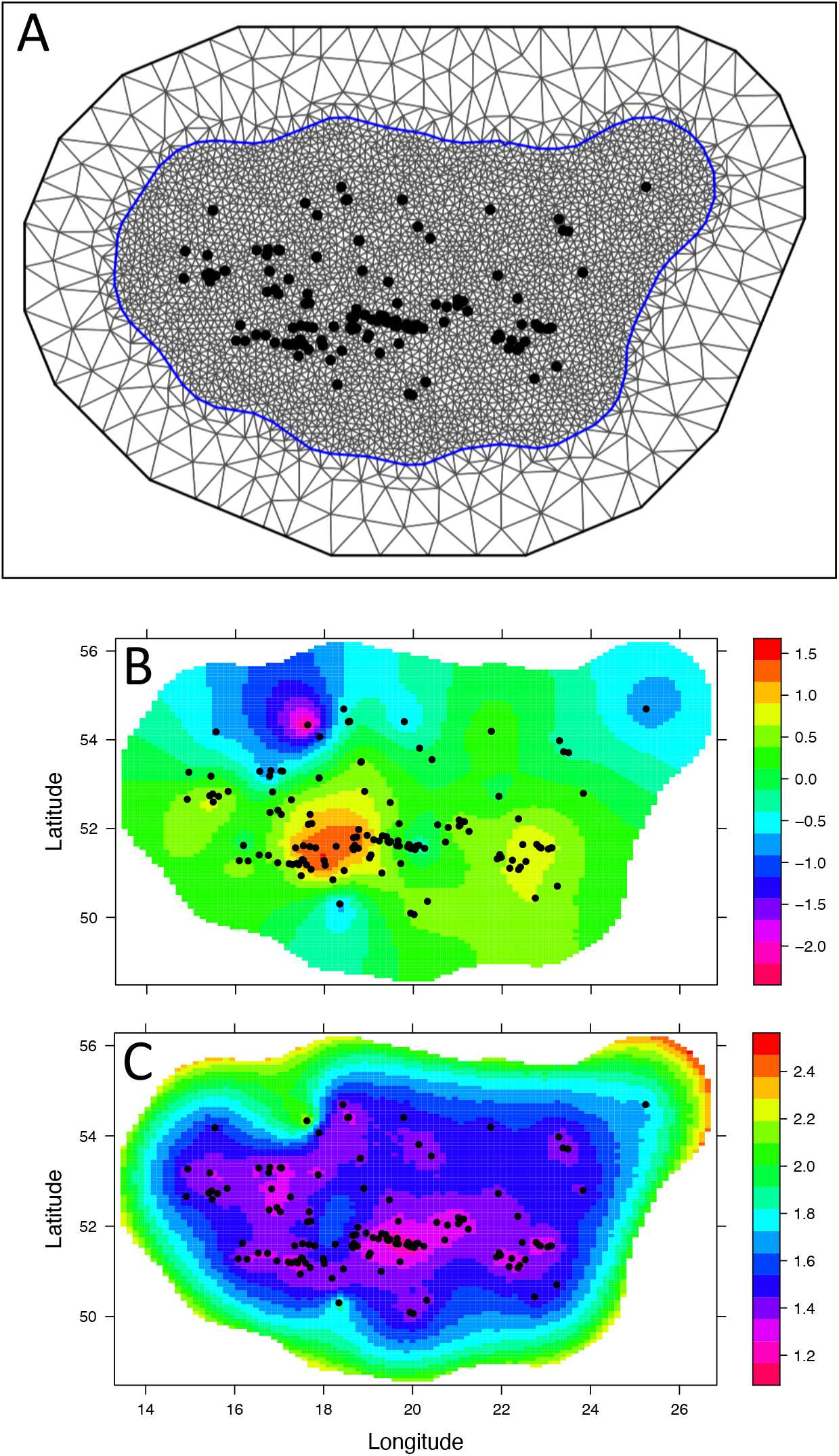
(A) Mesh of 3,874 vertices generated by constrained refined Delaunay triangulation; (B) Posterior mean values of the GMRF estimated for threespine stickleback plate number across Poland; (C) Variation (SD) of the GMRF estimated for threespine stickleback plate number.

In the best-fitting model, both body size and T_10_ were statistically important predictors of lateral plate number (Table 2, Fig 4A-C). As predicted, T_10_ showed a negative association with lateral plate number (Fig. 5A), while body size showed a positive association (Fig. 5B). Approximately 27% of the variation in lateral plate number was explained in the spatial random field and the GMRF indicated the strongest negative effects on plate number in the north-west and north-east of Poland and positive effects in central Poland. These locations also showed elevated variance of the GMRF (Fig. 3C). Spatial dependency in lateral plate number was important at a scale of 3.6 km (Table 2, Fig. 4D).

**Table 2.** Fixed effects from the best-fitting model for threespine stickleback lateral plate number in Poland. Credible intervals are the 2.5% and 97.5% quantiles. Hyperparameters are: κ_spatial_ = 1.78, σ_spatial_ = 1.31; range = 3.56 km. Credible intervals that do not include zero indicate statistical importance.

| Term | Posterior mean | 2.5% quantile | 97.5% quantile |
| --- | --- | --- | --- |
| Intercept | 28.77 | 26.17 | 30.63 |
| Standard length (SL) | 1.07 | 0.53 | 1.60 |
| 10-year antecedent temperature ($T_{10}$) | -1.50 | -2.15 | -0.88 |

**Fig. 4.**
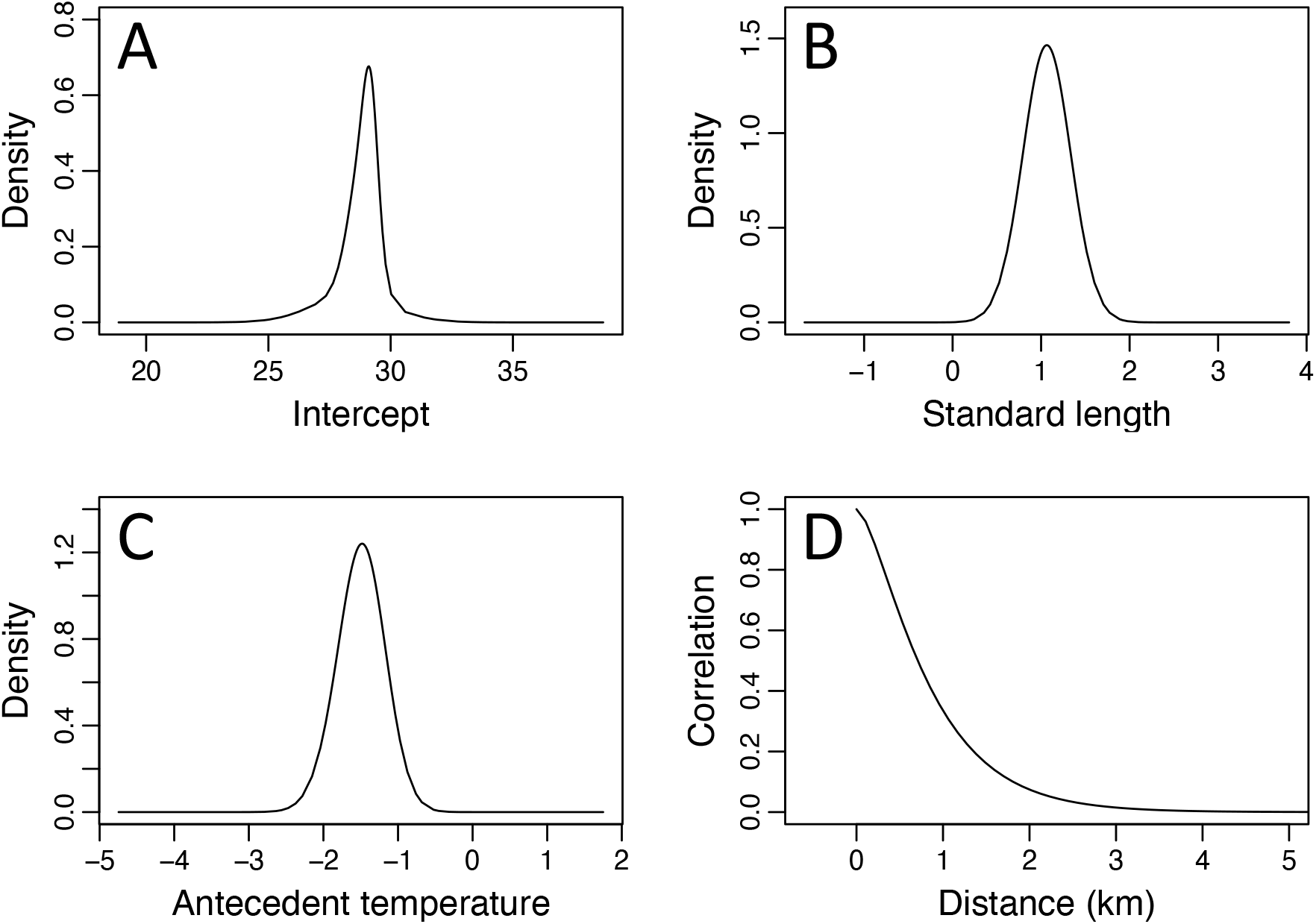
Posterior (marginal) distributions for: (A) model intercept; (B) standard length; (C) average 10-year antecedent air temperature; (D) Matérn correlation function.

**Fig. 5.**
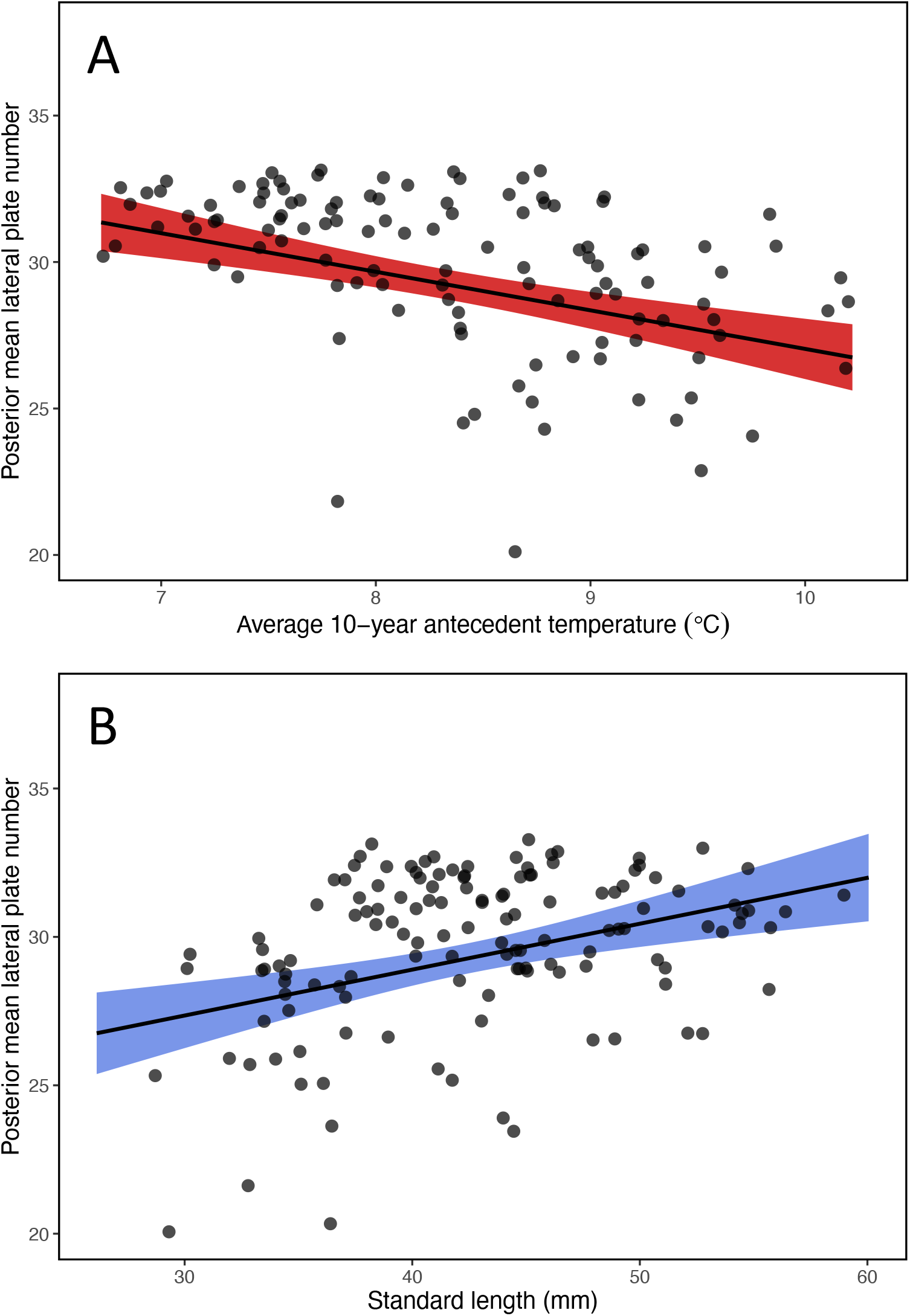
Posterior mean fitted estimates of lateral plate number for threespine sticklebacks as a function of: (A) average 10-year antecedent air temperature; (B) population standard length (mm). Shaded areas are 95% credible intervals. Black points are observed data for stickleback populations.

## Discussion

Using a latent Gaussian model, we provide evidence that over multiple decades, and on a large spatial scale, variation in threespine stickleback lateral plate number is predicted by antecedent air temperature. This effect is mediated by body size; elevated temperatures select for reduced body size in the threespine stickleback, which is negatively correlated with plate number (Fig. 5; Smith et al. 2020b). Strong negative effects of the GMRF on plate number were observed in the north-west and north-east of Poland (Fig. 3B), with both locations also showing elevated variance (Fig. 3C). In the north-west of Poland, the River Oder debouches into the Szczecin Lagoon and the Baltic Sea, permitting heavily-plated *complete* ecomorph threespine sticklebacks to penetrate freshwater systems from the Baltic. In the north-east of the area for which data were available (now in south-east Lithuania) minimum winter temperatures are extremely low for the region, sometimes falling below −30 °C. Thus, these effects are proposed as being a function of large-bodied and heavily-armored sticklebacks penetrating inland through a large river and lagoon system, in combination with selection for large body size in a region of unusually low environmental temperatures. Strong positive effects of the GMRF were also observed in Central Poland. In this region, plate numbers are depressed (Fig. 3B) and with low variance (Fig. 3C). This pattern may reflect local thermal pollution associated with the discharge of heated waste water from large cities, such as Warsaw and Łódź (Kalinowska et al., 2012).

The findings of this study are significant in demonstrating that temporal change in temperature can drive rapid natural adaptation in a phenotypic trait with a known genetic basis. Many studies that show phenotypic responses to climate change represent modifications to highly plastic traits, such as fecundity, survival, timing of migration and reproductive phenology (Crozier and Hutchings 2014). Growth is also a plastic trait in the threespine stickleback (e.g. Mccairns and Bernatchez 2012), though plate morphotype (Colosimo et al. 2005; Barrett et al. 2008; Marchinko 2009) and number (Hansson et al. 2016) are not. The effect demonstrated here represents a rare example of a phenotypic response to temperature change that is underpinned by genetic change. Importantly, the result does not simply represent variance in phenotype along a spatial climatic gradient, which can be a reflection of adaptation to contrasting environmental conditions. Instead, after controlling for spatial patterns in the data, we show a broad trend of declining plate number in response to a long-term increase in environmental temperature.

An association between threespine stickleback lateral plate number and temperature has been recognized for several decades, though it has never been adequately explained (Heuts, 1947; Hagen & Moodie, 1982; Wootton, 2009). High numbers of lateral plates are particularly characteristic of fresh water populations exposed to low winter temperatures on the eastern and northern fringes of continents (Wootton, 1976; Hagen & Moodie, 1982). In contrast, low numbers of lateral plates are associated with mild winter temperatures in the south and west of the geographic range of the species (Wootton 2009; Smith et al. 2020b). Despite this striking pattern, the association between plate number and temperature has not been a feature of discussions on other potential selective agents for the evolution of plate number in the threespine stickleback, at least until recently (Des Roches et al. 2019; Smith et al. 2020b).

It is unclear whether reduced plate number represents an adaptive response to elevated temperatures or a correlated response to selection for reduced body size. If a reduction in plate number is mediated by a change in body size, as proposed here, there may be no adaptive advantage to fewer lateral plates at elevated temperatures. Reduced body size as a response to higher temperatures is a prediction of Bergmann’s rule (Bergmann, 1847). The causal explanation for Bergmann’s rule is unclear but applies across a range of endotherms and ectotherms, including fishes (Belk & Houston, 2002). In the case of ectotherms, higher resting metabolic rates may incur higher energetic costs that compromise growth. Selection for large offspring body size at low temperatures may also play a role (Pettersen et al., 2019).

An implication of a correlated response between body size and lateral plate number is that variation in lateral plate number has no Darwinian adaptive advantage and solely reflects a scaling relationship *sensu* Thompson (1917). Alternatively, natural selection may drive changes in body size in response to temperature, with plate number varying to optimize mechanical efficiency at a given body size (Bonner & Horn, 2000). In aquatic animals scaling effects have particular consequences for hydrodynamic resistance to movement (Schmidt-Nielsen, 1984). Lateral plates increase drag in the threespine stickleback and correlate negatively with swimming velocity after correcting for body size (Bergstrom, 2002). An outcome is that selection on sticklebacks for armor loss will minimize drag (Walker, 1997; Bergstrom, 2002), and sticklebacks that experience selection for reduced body size at elevated temperatures are predicted to simultaneously experience selection for plate loss to avoid compromising hydrodynamic efficiency (Smith et al., 2020b).

The threespine stickleback is common and widespread species across coastal regions of the northern hemisphere (Wootton 1976). It is a vertebrate model in evolutionary, genomic, behavioral and ecological studies, and has been the subject of a broad range of research questions for over a century. Lateral plate phenotype offers a potential tool for monitoring environmental change in historical data sets and archived samples, and potentially also in the fossil record (Bell 2009). This study demonstrates the utility of the threespine stickleback as a model species for measuring the evolutionary and ecological impacts of climate change across the northern hemisphere.

## Acknowledgements

We are grateful to Jerzy Bańbura, Martin Reichard and Rowena Spence for comments. Research was supported by the POLONEZ Fellowship of National Science Centre, Poland (2015/19/P/NZ8/03582). This project has received funding from the European Union’s Horizon 2020 research and innovation program under the Marie Skłodowska-Curie grant agreement No. 665778.

