## Supplemental Table 1 for "Elevated temperatures drive the evolution of armor loss in the threespine stickleback *Gasterosteus aculeatus*"

**Table S1.** Location of collection of threespine stickleback samples used in the study, showing nearest meteorological station, year of collection and source. 'Collected' indicates samples collected by the authors in the course of the study, and 'Institute of Zoology' indicates samples from the Museum of the Institute of Zoology of the Polish Academy of Sciences in Warsaw.

| Site | Code | Latitude | Longitude | Station | Year | Source |
|---|---|---|---|---|---|---|
| towa | 2 | 51.528119 | 20.039665 | 922 | 1947 | Penczak (1960) |
| wik2 | 3 | 53.139844 | 17.878421 | 922 | 1954 | Penczak (1960) |
| alow | 4 | 51.838172 | 19.307270 | 922 | 1955 | Penczak (1960) |
| jaow | 5 | 52.084612 | 20.546161 | 922 | 1955 | Penczak (1960) |
| jeka | 6 | 52.155549 | 21.113025 | 922 | 1955 | Penczak (1960) |
| olka | 7 | 51.710796 | 19.488621 | 922 | 1955 | Penczak (1960) |
| rina | 8 | 52.414744 | 16.951666 | 922 | 1955 | Penczak (1960) |
| bycz | 9 | 53.139849 | 17.878422 | 922 | 1955 | Bańbura (1988) |
| hel2 | 10 | 54.601241 | 18.802503 | 779 | 1957 | Penczak (1962) |
| neer | 11 | 51.708859 | 19.426277 | 922 | 1957 | Penczak (1960) |
| biza | 12 | 52.793106 | 23.828180 | 922 | 1958 | Bańbura (1988) |
| kuce | 13 | 54.735396 | 18.576306 | 779 | 1959 | Bańbura (1988) |
| grdz | 14 | 53.505324 | 18.830658 | 779 | 1959 | Bańbura (1988) |
| trka | 15 | 53.495690 | 18.817316 | 779 | 1959 | Bańbura (1988) |
| nazw | 16 | 51.287623 | 22.230568 | 922 | 1960 | Bańbura (1988) |
| zaec | 17 | 52.077261 | 21.050635 | 209 | 1961 | Bańbura (1988) |
| slow | 18 | 50.360263 | 20.318465 | 209 | 1961 | Bańbura (1988) |
| swna | 19 | 53.921852 | 14.279426 | 208 | 1961 | Bańbura (1988) |
| waca | 20 | 53.890727 | 14.415257 | 208 | 1961 | Bańbura (1988) |
| zasc | 21 | 50.703458 | 23.242101 | 922 | 1961 | Penczak (1962) |
| ruka | 22 | 53.180449 | 15.443586 | 208 | 1962 | Penczak (1962) |
| kana | 23 | 54.190520 | 21.756626 | 205 | 1962 | Bańbura (1988) |
| frki | 24 | 53.979806 | 23.289989 | 2024 | 1972 | Bańbura (1988) |
| scwa | 25 | 51.396473 | 16.739423 | 210 | 1975 | Bańbura (1988) |
| liko | 26 | 53.730893 | 23.383218 | 2024 | 1979 | Bańbura (1988) |
| bora | 27 | 53.709828 | 23.496416 | 2024 | 1980 | Bańbura (1988) |
| mlow | 28 | 52.024634 | 20.785172 | 209 | 1981 | Bańbura (1988) |
| luwa | 29 | 51.568152 | 20.050315 | 926 | 1984 | Bańbura (1988) |
| wlo2 | 30 | 54.787399 | 18.423112 | 205 | 1984 | Bańbura (1988) |
| goka | 31 | 51.847170 | 18.958908 | 206 | 1985 | Bańbura (1988) |
| soca | 32 | 51.539252 | 23.038501 | 927 | 1985 | Bańbura (1988) |
| laa2 | 33 | 51.551808 | 19.952287 | 926 | 1985 | Bańbura (1988) |
| laa1 | 34 | 51.554482 | 19.940110 | 926 | 1985 | Bańbura (1988) |
| boki | 35 | 51.563831 | 22.874985 | 927 | 1985 | Bańbura (1988) |
| wyno | 36 | 51.628623 | 19.913480 | 926 | 1985 | Bańbura (1988) |
| zaow | 37 | 51.560802 | 19.960158 | 926 | 1985 | Bańbura (1988) |
| szra | 38 | 51.206642 | 17.204977 | 210 | 1985 | Bańbura (1988) |
| dzow | 39 | 54.030975 | 14.766814 | 696 | 1985 | Bańbura (1988) |
| gdia | 40 | 54.515166 | 18.552570 | 205 | 1985 | Bańbura (1988) |

| | | | | | | |
|---|---|---|---|---|---|---|
| stca | 41 | 53.297405 | 17.023680 | 2026 | 1985 | Bańbura (1988) |
| szki | 42 | 51.271306 | 16.280638 | 210 | 1985 | Bańbura (1988) |
| kog2 | 43 | 54.178284 | 15.559872 | 2022 | 1985 | Bańbura (1988) |
| kog1 | 44 | 54.187556 | 15.555853 | 2022 | 1985 | Bańbura (1988) |
| scow | 45 | 51.339282 | 19.014996 | 924 | 1985 | Bańbura (1988) |
| trpa | 46 | 51.211504 | 19.717678 | 926 | 1985 | Bańbura (1988) |
| luta | 47 | 53.809496 | 20.140522 | 752 | 1985 | Bańbura (1988) |
| dzin | 48 | 51.620491 | 18.680373 | 924 | 1985 | Bańbura (1988) |
| miwo | 49 | 54.358462 | 18.947591 | 205 | 1985 | Bańbura (1988) |
| sota | 50 | 52.110142 | 19.681124 | 206 | 1985 | Bańbura (1988) |
| mrno | 51 | 54.149346 | 15.302109 | 2022 | 1985 | Bańbura (1988) |
| chia | 52 | 51.563999 | 18.636448 | 924 | 1985 | Bańbura (1988) |
| stka | 53 | 51.682039 | 19.459032 | 926 | 1985 | Bańbura (1988) |
| olwa | 54 | 54.404609 | 18.536422 | 205 | 1985 | Bańbura (1988) |
| peew | 55 | 51.810011 | 18.727500 | 206 | 1985 | Bańbura (1988) |
| czla | 56 | 51.749185 | 19.109527 | 206 | 1985 | Bańbura (1988) |
| moa2 | 57 | 51.565899 | 23.121402 | 927 | 1985 | Bańbura (1988) |
| grin | 58 | 51.277857 | 16.081216 | 210 | 1985 | Bańbura (1988) |
| buow | 59 | 51.581272 | 22.840616 | 927 | 1985 | Bańbura (1988) |
| sien | 60 | 51.643188 | 22.778235 | 927 | 1985 | Bańbura (1988) |
| moa1 | 61 | 51.564823 | 23.099045 | 927 | 1985 | Bańbura (1988) |
| wlo1 | 62 | 54.796619 | 18.415043 | 205 | 1985 | Bańbura (1988) |
| czin | 63 | 51.605453 | 19.684552 | 926 | 1985 | Bańbura (1988) |
| deka | 64 | 51.526809 | 18.657447 | 924 | 1985 | Bańbura (1988) |
| sidz | 65 | 51.556382 | 18.761338 | 924 | 1985 | Bańbura (1988) |
| deca | 66 | 51.566356 | 17.800683 | 924 | 1986 | Bańbura (1988) |
| suce | 67 | 51.613862 | 17.525792 | 210 | 1986 | Bańbura (1988) |
| ryol | 68 | 52.825357 | 16.830538 | 2029 | 1986 | Bańbura (1988) |
| odow | 69 | 51.591885 | 17.670582 | 924 | 1986 | Bańbura (1988) |
| maew | 70 | 50.433980 | 22.745754 | 207 | 1986 | Bańbura (1988) |
| gowo | 71 | 52.724528 | 15.615780 | 2029 | 1986 | Bańbura (1988) |
| ola1 | 72 | 51.214936 | 17.376330 | 210 | 1986 | Bańbura (1988) |
| brny | 73 | 51.598138 | 18.262501 | 924 | 1986 | Bańbura (1988) |
| micz | 74 | 51.566657 | 17.348729 | 210 | 1986 | Bańbura (1988) |
| drko | 75 | 52.837147 | 15.831417 | 2029 | 1986 | Bańbura (1988) |
| jeko | 76 | 51.795562 | 18.654289 | 206 | 1986 | Bańbura (1988) |
| gika | 77 | 54.693479 | 18.432708 | 205 | 1987 | Bańbura (1988) |
| hel1 | 78 | 54.601241 | 18.802503 | 205 | 1987 | Bańbura (1988) |
| weki | 79 | 52.319653 | 17.675142 | 206 | 1987 | Bańbura (1988) |
| gesh | 80 | 54.409683 | 18.560876 | 205 | 2001 | Collected |
| vila | 81 | 51.935231 | 21.254267 | 209 | 2006 | Collected |
| naow | 82 | 51.352009 | 21.980463 | 333 | 2011 | Collected |
| czje | 83 | 53.291620 | 17.057621 | 2026 | 2013 | Collected |
| doca | 84 | 53.288572 | 16.543036 | 2026 | 2013 | Collected |
| glia | 85 | 50.303603 | 18.342619 | 928 | 2013 | Collected |
| gwda | 86 | 53.181727 | 16.766054 | 2026 | 2013 | Collected |
| maka | 87 | 53.553937 | 20.420164 | 752 | 2013 | Collected |

| | | | | | | |
|---|---|---|---|---|---|---|
| bina | 88 | 51.551988 | 19.951603 | 926 | 2017 | Collected |
| gaac | 89 | 51.622267 | 20.124388 | 926 | 2017 | Collected |
| grna | 90 | 51.405483 | 19.046740 | 924 | 2017 | Collected |
| lusk | 91 | 51.720441 | 19.234491 | 926 | 2017 | Collected |
| mala | 92 | 52.648323 | 17.255117 | 2027 | 2017 | Collected |
| mige | 93 | 51.727531 | 19.655385 | 926 | 2017 | Collected |
| mlka | 94 | 52.094357 | 17.637058 | 206 | 2017 | Collected |
| muka | 95 | 51.696023 | 20.725335 | 209 | 2017 | Collected |
| paka | 96 | 51.588021 | 19.870486 | 926 | 2017 | Collected |
| pice | 97 | 51.641090 | 19.900667 | 926 | 2017 | Collected |
| plca | 98 | 53.304241 | 16.792910 | 2026 | 2017 | Collected |
| prik | 99 | 50.094560 | 19.940044 | 929 | 2017 | Collected |
| prna | 100 | 51.054472 | 18.443136 | 924 | 2017 | Collected |
| runa | 101 | 51.620696 | 16.176834 | 923 | 2017 | Collected |
| sana | 102 | 52.780747 | 15.483672 | 2029 | 2017 | Collected |
| stec | 103 | 52.737913 | 15.417737 | 2029 | 2017 | Collected |
| toor | 104 | 51.191223 | 17.279330 | 210 | 2017 | Collected |
| waa2 | 105 | 52.598476 | 15.495014 | 2029 | 2017 | Collected |
| waa1 | 106 | 51.981048 | 18.770877 | 206 | 2017 | Collected |
| woka | 107 | 51.614571 | 19.625192 | 926 | 2017 | Collected |
| basz | 108 | 52.111908 | 17.714376 | 206 | 2018 | Collected |
| buka | 109 | 50.846994 | 18.189440 | 924 | 2018 | Collected |
| chka | 110 | 51.554514 | 20.255729 | 926 | 2018 | Collected |
| czka | 111 | 51.258106 | 22.533083 | 927 | 2018 | Collected |
| doka | 112 | 50.939189 | 17.475575 | 210 | 2018 | Collected |
| doyw | 113 | 51.183600 | 17.595830 | 210 | 2018 | Collected |
| goca | 114 | 53.268085 | 14.946573 | 208 | 2018 | Collected |
| jaca | 115 | 51.174445 | 18.015636 | 924 | 2018 | Collected |
| jyka | 116 | 51.405841 | 16.530701 | 210 | 2018 | Collected |
| kock | 117 | 51.638954 | 22.463388 | 333 | 2018 | Collected |
| kola | 118 | 52.318293 | 17.026521 | 923 | 2018 | Collected |
| krka | 119 | 51.106924 | 22.170501 | 927 | 2018 | Collected |
| lawa | 120 | 51.230914 | 16.947981 | 210 | 2018 | Collected |
| luia | 121 | 54.334398 | 17.623947 | 332 | 2018 | Collected |
| neca | 122 | 51.134008 | 22.411787 | 927 | 2018 | Collected |
| niob | 123 | 51.280252 | 17.992182 | 924 | 2018 | Collected |
| ola2 | 124 | 51.291757 | 17.509940 | 210 | 2018 | Collected |
| olki | 125 | 54.412791 | 18.567814 | 205 | 2018 | Collected |
| olza | 126 | 51.606304 | 20.065285 | 926 | 2018 | Collected |
| osna | 127 | 51.286618 | 17.522615 | 210 | 2018 | Collected |
| rika | 129 | 51.818148 | 19.395414 | 926 | 2018 | Collected |
| skka | 130 | 51.076659 | 22.370395 | 927 | 2018 | Collected |
| soka | 131 | 51.296547 | 17.495735 | 210 | 2018 | Collected |
| stga | 132 | 52.725120 | 21.926683 | 333 | 2018 | Collected |
| swza | 133 | 51.185978 | 17.420466 | 210 | 2018 | Collected |
| szow | 134 | 51.286605 | 17.522283 | 210 | 2018 | Collected |
| trca | 135 | 51.174588 | 18.015393 | 924 | 2018 | Collected |

| | | | | | | |
|---|---|---|---|---|---|---|
| trha | 136 | 54.066701 | 17.895552 | 2026 | 2018 | Collected |
| wiwa | 137 | 51.082118 | 17.698533 | 210 | 2018 | Collected |
| wiow | 138 | 51.000121 | 19.292859 | 926 | 2018 | Collected |
| wika | 139 | 52.363957 | 16.774023 | 923 | 2018 | Collected |
| wila | 140 | 52.585190 | 19.479726 | 206 | 2018 | Collected |
| wina | 141 | 52.658830 | 14.911167 | 2029 | 2018 | Collected |
| niwa | 224 | 52.836600 | 18.904057 | 779 | 1959 | Institute of Zoology |
| zaec | 225 | 52.056692 | 21.030120 | 922 | 1953 | Institute of Zoology |
| pace | 226 | 51.648695 | 19.369178 | NA | 1936 | Institute of Zoology |
| puwy | 227 | 51.414226 | 21.947346 | 922 | 1947 | Institute of Zoology |
| pasa | 228 | 54.408005 | 19.797314 | 205 | 1963 | Institute of Zoology |
| wima | 229 | 52.155215 | 21.149927 | 2022 | 1930 | Institute of Zoology |
| wino | 230 | 52.192681 | 21.030105 | 2022 | 1927 | Institute of Zoology |
| ldan | 231 | 51.582495 | 19.240398 | NA | 1936 | Institute of Zoology |
| jaec | 232 | 51.316921 | 21.889444 | 2022 | 1929 | Institute of Zoology |
| czny | 233 | 50.063406 | 20.016739 | 208 | 1930 | Institute of Zoology |
| laec | 234 | 52.952836 | 22.293832 | NA | 1932 | Institute of Zoology |
| liec | 235 | 52.218722 | 22.366589 | 2022 | 1930 | Institute of Zoology |
| wilo | 236 | 54.694441 | 25.240666 | 2022 | 1928 | Institute of Zoology |
| stun | 237 | 53.867948 | 23.106441 | NA | 1933 | Institute of Zoology |
| haza | 238 | 54.120054 | 22.912508 | NA | 1932 | Institute of Zoology |
