## Supplemental Table 2 for "Elevated temperatures drive the evolution of armor loss in the threespine stickleback *Gasterosteus aculeatus*"

**Table S2.** Meteorological stations for which data were used to estimate 10-year antecedent air temperatures for sampling locations at which threespine sticklebacks were collected. Daily temperature data for these sites are available from the European Climate Assessment & Dataset project at: www.ecad.eu (Klein Tank et al. 2002).

| Location | Code | Latitude | Longitude | Start date |
|---|---|---|---|---|
| VILNIUS | 202 | 54.633148 | 25.101128 | 1881 |
| BIALYSTOK | 204 | 53.106936 | 23.161954 | 1963 |
| HEL | 205 | 54.602753 | 18.813069 | 1951 |
| POZNAN | 206 | 52.199753 | 18.660204 | 1951 |
| RZESZOW JASIONKA | 207 | 50.115168 | 22.024515 | 1952 |
| SZCZECIN | 208 | 53.395277 | 14.622707 | 1920 |
| WARSZAWA-OKECIE | 209 | 52.162862 | 20.961142 | 1951 |
| WROCLAW | 210 | 51.103175 | 16.899849 | 1951 |
| LEBA | 332 | 54.753654 | 17.534747 | 1961 |
| SIEDLCE | 333 | 52.181003 | 22.24476 | 1966 |
| ELBLAG | 396 | 54.223192 | 19.543505 | 1961 |
| SWINOUJSCIE | 696 | 53.923359 | 14.242268 | 1961 |
| OLSZTYN | 752 | 53.768562 | 20.421349 | 1966 |
| SLUBICE | 779 | 52.348531 | 14.619554 | 1947 |
| TERESPOL | 922 | 52.078672 | 23.621987 | 1944 |
| LESZNO | 923 | 51.835541 | 16.534707 | 1966 |
| WIELUN | 924 | 51.210222 | 18.556577 | 1966 |
| SULEJOW | 926 | 51.353316 | 19.86637 | 1966 |
| WLODAWA | 927 | 51.553556 | 23.529607 | 1966 |
| RACIBORZ | 928 | 50.061024 | 18.190816 | 1966 |
| KRAKOW | 929 | 50.074784 | 19.795591 | 1966 |
| BIELSKO-BIALA | 930 | 49.806381 | 19.000888 | 1966 |
| NOWY SACZ | 932 | 49.627162 | 20.688642 | 1966 |
| LESKO | 933 | 49.466466 | 22.341772 | 1966 |
| KOSZALIN | 2022 | 54.204519 | 16.155148 | 1920 |
| SUWALKI | 2024 | 54.130836 | 22.948926 | 1961 |
| CHOJNICE | 2026 | 53.715049 | 17.531601 | 1966 |
| TORUN | 2027 | 53.042098 | 18.595232 | 1966 |
| MLAWA | 2028 | 53.104193 | 20.360955 | 1966 |
| GORZOW WLKP | 2029 | 52.740755 | 15.277372 | 1966 |
| ZIELONA GORA | 2031 | 51.929968 | 15.524727 | 1966 |
| KLODZKO | 2036 | 50.436872 | 16.614213 | 1966 |
| KATOWICE | 2037 | 50.241002 | 19.032145 | 1966 |
| KIELCE | 2038 | 50.810477 | 20.692247 | 1966 |
